# Predictability of Autism, Schizophrenic and Obsessive Spectra Diagnosis: Toward a Damage Network Approach

**DOI:** 10.1101/014563

**Authors:** Franco Cauda, Tommaso Costa, Luciamo Fava, Sara Palermo, Francesca Bianco, Sergio Duca, Giuliano Geminiani, Karina Tatu, Roberto Keller

## Abstract

Schizophrenia, obsessive-compulsive and autistic disorders are traditionally considered as three separate psychiatric conditions each with specific symptoms and pattern of brain alterations. This view can be challenged since these three conditions have the same neurobiological origin, stemming from a common root of a unique neurodevelopmental tree.

The aim of this meta-analytic study was to determine, from a neuroimaging perspective, whether i) white matter and gray matter alterations are specific for the three different spectrum disorders, and the nosographical differentiation of three spectra is supported by different patterns of brain alterations. ii) it might be possible to define new spectra starting from specific brain damage. iii) it is possible to detect a “brain damage network” (a connecting link between the damaged areas that relates areas constantly involved in the disorder).

Three main findings emerged from our meta-analysis:

1. The three psychiatric spectra do not appear to have their own specific damage.
2. It is possible to define two new damage clusters. The first includes substantial parts of the salience network, and the second is more closely linked to the auditory-visual, auditory and visual somatic areas.
3. It is possible to define a "Damage Network" and to infer a hierarchy of brain substrates in the pattern of propagation of the damage.

These results suggest the presence of a common pattern of damage in the three pathologies plus a series of variable alterations that, rather than support the sub-division into three spectra, highlight a two-cluster parcellation with an input-output and more cognitive clusters.

## INTRODUCTION

Ever since Kanner (Kanner, 1943) derived the term “autism” from Bleuer’s (Bleuler, 1911) description of schizophrenia to indicate the new syndrome in childhood, it has been recognized that there is a strict link between schizophrenia and autism. Moreover, the term childhood schizophrenia has encompassed the condition we now recognize as autism since DSM-III (American Psychiatric Association, 1980) clearly distinguished schizophrenia from autism and, furthermore, DSM-IV-TR (American Psychiatric Association, 2000) reported only a possible comorbidity between two distinct entities. Autism spectrum disorder (ASD) is now defined by: 1) persistent deficit in social communication and social interaction across contexts; 2) restricted, repetitive patterns of behavior, interest, or activities that include atypical sensory behavior. The symptoms that impair everyday functioning must be present in the early developmental period (but may not become fully manifest until social demands exceed limited capacities, or may be masked by learned strategies in later life) (American Psychiatric Association, 2013). Schizophrenia spectrum (SCZS) includes, in DSM-5 (American Psychiatric Association, 2013), schizophrenia, other psychotic disorders, and schizotypal (personality) disorder. A genetic basis for schizophrenia spectrum disorders has been validated using molecular polygene scores from genome-wide association studies (GWAS) (Bigdeli et al., 2014).

Even changing from a categorical to a spectrum model, diagnostic criteria separate the two spectra (American Psychiatric Association, 2013; World Health Organization, 1992).

Conversely, other authors, adopting a neurodevelopmental approach, have stated that these two spectra may instead reflect the influence of the level of maturation on the overt expression of a common single disease process (Cheung et al., 2010; de Lacy and King, 2013; King and Lord, 2011; Owen et al., 2011; Stone and Iguchi, 2011).

This intriguing debate could be explored by considering the fact that diagnoses are formulated phenomenologically rather than biologically, as epiphenomenic collections of symptoms and patterns of behavior, but since ASD and SCZS accumulate more and more evidence of epidemiologic, genetic, molecular, brain imaging, and also some behavioral levels of convergence (Cheung et al., 2010; de Lacy and King, 2013; King and Lord, 2011; Stone and Iguchi, 2011), it is not possible to hypothesize a common root. ASD and SCZS show a similar general prevalence (1%, higher in males), high comorbidity of up to 60% (Rapoport et al., 2009; Unenge Hallerback et al., 2012) and similar heritable endophenotypes such as deficits in Theory of Mind (de Achaval et al., 2010; Fett et al., 2011; Mehl et al., 2010; Williams and Happe, 2010) and in mirror neuron function (Boria et al., 2009; Enticott et al., 2008; Keller et al., 2011; Nagy et al., 2010; Rizzolatti and Craighero, 2004). There is also some genetic overlapping as in SHANK3 (Bourgeron, 2009; Bourgeron et al., 2009; Gauthier et al., 2010) or GWAS, candidate genes and copy number variants (CNVs) studies (Carroll and Owen, 2009; Poelmans et al., 2013; Rodriguez-Murillo et al., 2012). Further similarities between the two spectra concern epigenetic dysregulation (Leblond et al., 2012; McCarthy et al., 2014) and neuropathological abnormalities related to a disruption of early events in cerebral development such as migration, lamination and proliferative defects (de Lacy and King, 2013). Neuroimaging studies have also revealed some comparable regional gray and white matter abnormalities and altered patterns of structural and functional connectivity in both ASD and SCZS (Cauda et al., 2014; Cauda et al., 2011; Cheung et al., 2010; Mueller et al., 2012; Zalesky et al., 2011).

Other disorders showing an intriguing link to ASD are those related to the obsessive-compulsive spectrum disorder (OCDS), defined now in DSM-5 as obsessive-compulsive and related disorder (American Psychiatric Association, 2013). The restricted, repetitive patterns of behavior, interest, or activities and the excessive adherence to routines, ritualized patterns of verbal/non-verbal behavior, the excessive resistance to change and the highly restricted, fixated interests described in ASD closely resemble the symptoms observed in OCDS. A common neurobiology of repetitive behavior in ASD and OCDS has been underlined (Langen et al., 2011a; Langen et al., 2011b). Moreover, not only are ASD and SCZS linked, but there might also be a pathogenetic bridge between OCDS and SCZS (Owashi et al., 2010). OCD prevalence in SCZ is 13.6% and obsessive-compulsive symptoms in SCZ are present in 30.3% of cases (Swets et al., 2014). The presence of obsessive-compulsive symptoms in chronic schizophrenia is not only associated with greater cognitive impairment but also with increased difficulties with at least some aspects of social cognitive function (Whitton and Henry, 2013).

Given these premises, we decided to explore the possibility of unifying, or otherwise clearly distinguishing ASD from SCZS and from OCDS, based on the pattern of brain alterations, from a psychoneurobiological perspective. Indeed the first aim of this meta-analytic study was to determine, from a neuroimaging perspective, whether white matter (WM) and gray matter (GM) alterations are specific for the three different spectrum disorders, and the nosographical differentiation of the three spectra is supported by different patterns of brain alterations.

Second, we investigated the possibility of defining new spectra starting from specific brain damage, from a neuroimaging perspective.

Third we investigated the possibility of detecting a “brain damage network”, meaning a connecting link between the damaged areas that, very much like in a neurophysiological circuit, relates specific areas that are constantly neuropathologically involved in the disorder, so that if one area is involved in the damage a second specific one, linked to the first one, is also probably damaged. If such a link exists, certain areas, if damaged, might be more capable than others of triggering a domino-effect on secondary areas. To the best of our knowledge, this damage network hypothesis has never previously been investigated using neuroimaging in vivo techniques.

## MATERIALS AND METHODS

### SELECTION OF STUDIES

We adopted the definition of meta-analysis embraced by the Cochrane Collaboration (Green et al., 2008) and the “PRISMA Statement” international guidelines to ensure transparent and complete reporting of data selection (Liberati et al., 2009; Moher et al., 2009).

A systematic search strategy was used to identify relevant studies, published on or before 31 August 2012, across the online database most frequently used in the international literature (Medline database with PubMed literature search: http://www.ncbi.nlm.nih.gov/pubmed). In an initial phase we analyzed the cognitive phenomics of all the keywords relevant to our purposes, permitting patterns in the literature to be seen (see the supplemental online material for more details). As input we chose different sets of "terms" and a relevance threshold: terms were dropped if they did not have at least 5 co-occurrences with some other term. Specifically, for ASD, OCDS and SCZS and for the single disorders included in each spectrum, as defined in DSM-5 (American Psychiatric Association, 2013), the search algorithm was constructed matching for Diffusion Tensor Imaging (DTI), and Voxel Based Morphometry (VBM). Association measures were analyzed to get perspective on the biomedical research literature: co-occurrences of the selected terms were analyzed using the natural logarithm of the Jaccard co-occurrence index (ln J) (Jaccard, 1901), a skew coefficient the use of which is preferable when dealing with lists derived from observations in which the representativeness of the sample is not entirely certain. The ln J index is a commonly-used measure of co-occurrence in a set of documents and is employed to compare the similarity and diversity of sample sets, being lower when objects are more alike (see the supplemental online material for more details and the results of this analysis).

Accordingly, up until 31 August 2013, 684 papers had been indexed on PubMed with the selected search terms. We also searched the bibliographies of published meta-analyses and reviews to identify additional studies which were not included in the PubMed literature search database. Nonetheless this outcome ensured the presence of a sufficient number of articles from among which to select the studies of interest to carry out a meaningful meta-analysis.

All included articles were analyzed and any that did not meet the inclusion criteria were excluded. Indeed all articles were reviewed to establish (1) the existence of the healthy control group for the pathological sample; (2) that the results were reported in Talairach/ Tournoux or in Montreal Neurological Institute (MNI) coordinates; (3) that the studies reported cerebral structural changes, as assessed by DTI or VBM; (4) that they were original works. We also tried to identify any instances of multiple reports of single data sets across articles to ensure that only one report of a study contributed to the coordinates for the present meta-analysis (See Graph 1: PRISMA Flow Diagram).

The studies were independently ascertained and checked by the authors for any discrepancies, which were resolved in a discussion phase. Descriptive information was extracted from each article. Based on these criteria, 139 papers were included in the analysis (see the Supplemental Online Material to the complete references), with an overall sample of 4719 subjects: 820 individuals within the ASD group, 661 in the OCDS group, and 3238 in the SCZS group. For a summary description of the papers selected and the selection procedure see Table 1 and the supplemental online material.

**Table 1.**
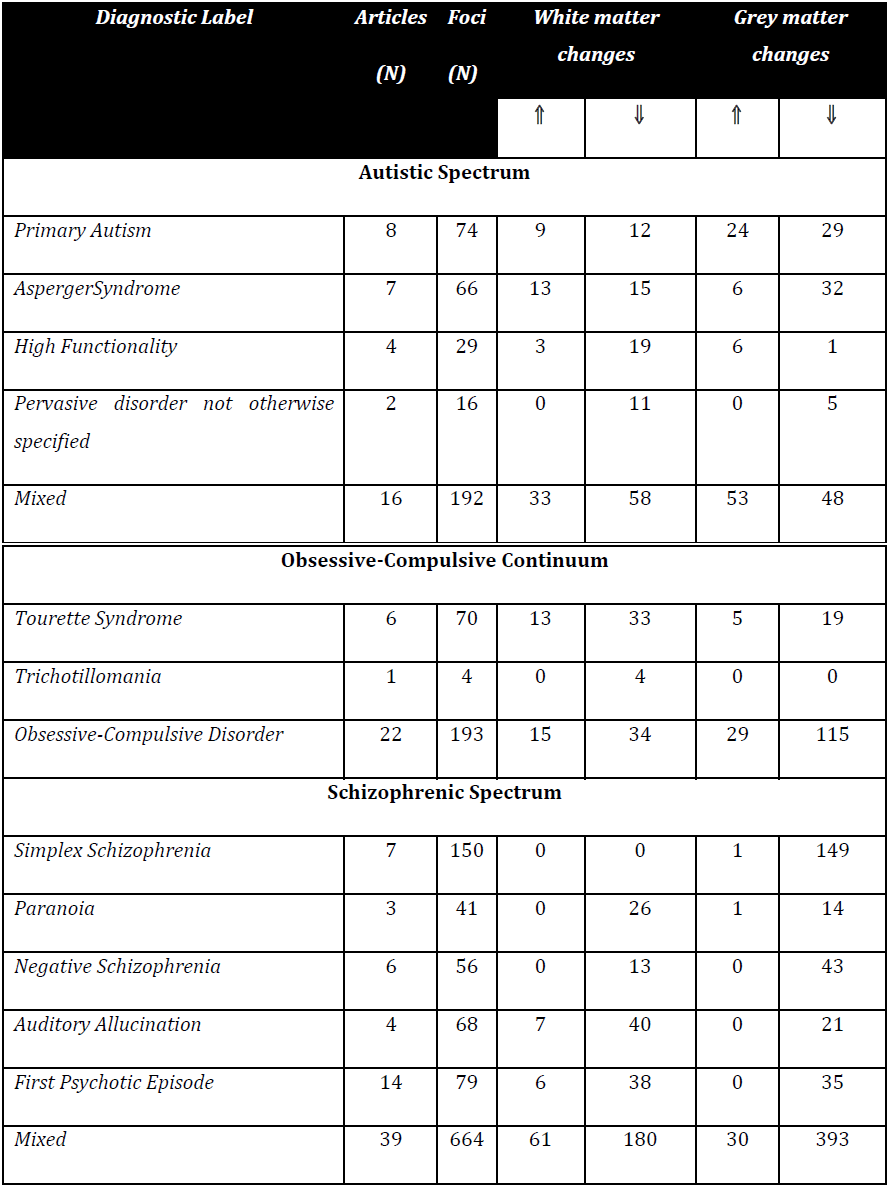
WM and GM variations with relative numbers of foci for each of the selected psychiatric spectrum.

| Diagnostic Label | Articles<br>(N) | Foci<br>(N) | White matter<br>changes |  | Grey matter<br>changes |  |
| --- | --- | --- | --- | --- | --- | --- |
|  |  |  | ↑ | ↓ | ↑ | ↓ |
|  |  |  | Autistic Spectrum |  |  |  |
| Primary Autism | 8 | 74 | 9 | 12 | 24 | 29 |
| AspergerSyndrome | 7 | 66 | 13 | 15 | 6 | 32 |
| High Functionality | 4 | 29 | 3 | 19 | 6 | 1 |
| Pervasive disorder not otherwise specified | 2 | 16 | 0 | 11 | 0 | 5 |
| Mixed | 16 | 192 | 33 | 58 | 53 | 48 |

| **Obsessive-Compulsive Continuum** |  |  |  |  |  |  |
| --- | --- | --- | --- | --- | --- | --- |
| *Tourette Syndrome* | 6 | 70 | 13 | 33 | 5 | 19 |
| *Trichotillomania* | 1 | 4 | 0 | 4 | 0 | 0 |
| *Obsessive-Compulsive Disorder* | 22 | 193 | 15 | 34 | 29 | 115 |
| **Schizophrenic Spectrum** |  |  |  |  |  |  |
| *Simplex Schizophrenia* | 7 | 150 | 0 | 0 | 1 | 149 |
| *Paranoia* | 3 | 41 | 0 | 26 | 1 | 14 |
| *Negative Schizophrenia* | 6 | 56 | 0 | 13 | 0 | 43 |
| *Auditory Allucination* | 4 | 68 | 7 | 40 | 0 | 21 |
| *First Psychotic Episode* | 14 | 79 | 6 | 38 | 0 | 35 |
| *Mixed* | 39 | 664 | 61 | 180 | 30 | 393 |

### ANATOMICAL LIKELIHOOD ESTIMATION AND MODELED ACTIVATION CREATION

We performed an Anatomical Likelihood Estimation (ALE) to statistically summarize the results of the experiments utilized. The ALE is a quantitative voxel-based meta-analysis technique able to give information about the anatomical reliability of results through a comparison using a sample of reference studies from the existing literature.

ALE meta-analysis is an appropriate tool for estimating consistent activations across several neuroimaging studies (Laird et al., 2005). This method is unable to use the foci coordinates of interest directly measured in the frame supplied by the authors; these must first be transformed into a secondary standard stereotactic space (Talairach standard coordinates).

A given meta-analysis considers each focus of each paper as the central point of a 3-dimensional Gaussian probability distribution; more in detail the Talairach space is subdivided into 2 mm^3^ volumes and the following probability density function (product of the three one-dimensional probability densities) is used:

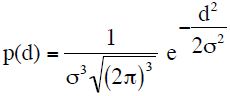

where d is the Euclidean distance between the voxels and the focus taken into account and σ is the standard deviation of the one-dimensional distribution. The standard deviation σ is easily obtained through the Full Width Half Maximum (FWHM) as:

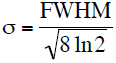

These Gaussian functions are then added to create a statistical map that estimates the likelihood of activation for each voxel determined by the whole set of studies. Furthermore, a threshold is fixed in the map using a permutation test, as suggested by the Authors of ALE methodology (Laird et al., 2005; Lancaster et al., 2007; Lancaster et al., 2000) and reported in the existing literature concerning this topic (*www.brainmap.org*).

The activation maps are usually obtained by means of ROI (Regions Of Interest) for which multiple studies have shown peaks of statistical significance for activation loci. In order to reduce the typical variability of neuroimaging studies due to the sample and labs, it is possible to employ an algorithm that estimates the spatial incertitude for each focus, taking into account the possible differences between studies referring to diversity in the sample amplitude (Eickhoff et al., 2009).

An activation map estimation is created by the logical union of the probabilities calculated for each focus: each paper is thus associated with a Modeled Activation map (MA). Finally, all MA maps give the total AND of calculated probabilities. Here the map’s values are tested against a null hypothesis such as if all of the map’s values had been obtained at random.

A FDR (False Discovery Rate) method is employed to correct multiple comparisons and to set the threshold, defined as the expected amount of rejected false voxels over the total of rejections (i.e. false negatives). For the final result a threshold is settled and corrected by means of multiple comparisons.

### MULTI DIMENSIONAL SCALING

To investigate the similarity between the different spectra of psychiatric disorders (represented by the MA maps generated by the different papers analyzed in this study) we used the multidimensional scaling (MDS) method. MDS mainly consists of data proximity analysis techniques to check hidden structures. To this aim, each MA map for each paper was transformed into a series of vectors containing all the values of the original matrix. We then built a representational similarity matrix by computing the correlation between all r vectors and similarly to create the distance matrix (or representational dissimilarity matrix) defined as 1–r (Cauda et al., 2014; Kriegeskorte et al., 2008). The distance matrix was subjected to the multidimensional scaling analysis to obtain a geometrical representation of data deviation. MDS transforms a set of multidimensional points into a set of points within a smaller dimensional space using a projection, trying to maintain the distances between the points unchanged. Through this technique MA maps having a smaller Euclidean distance (first quartile) are represented with a straight connection. The smaller the Euclidean distance the greater the representational similarity.

### DAMAGE NETWORK (MORPHOMETRIC COALTERATION NETWORK)

In this paper we aimed to introduce a novel method to characterize the “importance” of each damaged area in the process of brain alteration after brain pathologies. This analysis, very much as in a game of dominoes, investigates whether, when an area “A” is damaged, it’s damage is statistically concatenated with damage to one or more other brain areas “B”, “C”, etc. Tracing a network of damage co-occurrences enables us to examine i) how damage to one brain area is statistically connected to other damaged areas and ii) which areas, if damaged, are coupled to a more extended pattern of alterations.

In order to study the “damage connectivity” between several areas, we created coactivation maps using a connection probability estimation employing the maximum likelihood considering the total ALE map of all MA maps, with a threshold of p<0.01.

These areas were then parcellated using an anatomical atlas. ROI were employed in order to parcel the MA maps and create series of binary casual variables composed using the following algorithm: if a voxel in the activation map for a certain area is active, the variable is fixed to 1 in the binary map, otherwise it is 0. Therefore the null hypothesis considering each pair of binary variables A and B, corresponding to activation of 2 ROI, is that the probability of activation of B does not depend on the observed value of A, while the alternative hypothesis affirms such dependence.

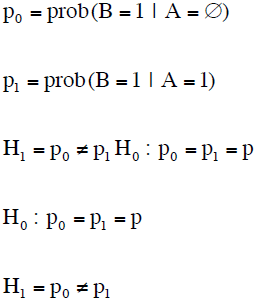

The consideration of these binary variable series affords the p value estimation as m/N, where m is the number of MA maps in which the ROI is active and N is the total number of MA maps. In a similar manner it is possible to define:

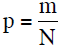

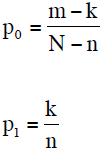

where n is the number of MA maps in which the ROI is active.

The likelihood-ratio test 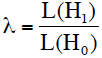 evaluates the alternative hypothesis H1 with respect to the null hypothesis H0. The λ distribution is shaped by a χ^2^ function with one freedom degree.

### NETWORK ANALYSIS TECNIQUES

To analyze the “Damage Network” we employed a network-based analysis technique. The analysis of complex networks is a new and powerful technique for quantifying the brain structure and the functional system. A network is defined as a system of nodes connected by a series of links. The topology of these networks may be described with a wide variety of measures.

### NODE DEGREE

The node degree is the number of connections that the node has to other nodes. We used the degree distribution to compare the node degree of different networks. Using the degree distribution it is possible to compare the network with a random network and the types of spectrum disorder. The degree distribution is the fraction of nodes with degree *k* defined as

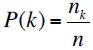

### NETWORK CLUSTERING

We employed the k-core decomposition algorithm (Alvarez-Hamelin et al., 2005; Bader and Hogue, 2003) to disentangle the hierarchical structures of our concordance network by progressively focusing on their central cores. The k-core decomposition of a network “N” works by recursively removing all the vertices of degree less than k, until all vertices in the remaining graph have at least degree k. This procedure allowed us to cluster our concordance network graph as its central most densely connected sub-graph.

### CLASSIFICATION

We used a machine learning approach to identify any distinguishing features of the different disorders. More specifically we used three different learning methods: the Naive Bayes classifier, the Support Vector Machines (SVM) and the k-nearest neighbors (knn).

The Naive Bayes classifier is based on Bayes’ theorem and assumes the independence of the features of each example. The maximum likelihood method is usually used in training sessions.

The support vector machine (SVM) is a supervised learning model with specific learning algorithms. The SVM constructs a hyper plane in a multidimensional space, with each axis representing a feature and a point in this space representing an example, used for the classification. A good separation is obtained if the hyper plane has the largest distance to the nearest data points of each class.

The k-nearest neighbor (knn) is a non-parametric method used in machine learning. In the learning phase, k is a constant, and an unlabeled vector is classified by assigning the label which is most frequent among the k training samples nearest to that point. The Euclidean distance is usually used as the continuous variable and a metric such as the Hamming distance for the discrete variable.

The performance of the various classifiers was evaluated using the Receiver operating characteristic (ROC) curve. This graphical plot is used to illustrate the performance of a binary classifier as a function of the discrimination threshold. The axis of the plot corresponds to the fraction of true positives out of the total positive named true positive rate (TPR) and to the fraction of false positives out of the total negative named false positive rate (FPR) at various thresholds. The TPR is also known as sensitivity and the FPR as fall-out.

In our case the input of the different classifiers was a vector of features (voxels) for each subject to form a matrix NxM with N subjects and M voxels and a vector representing the classes to which the subjects belong.

We used the leave-one-out method in which the learning algorithm is applied once for each instance, using all other instances as a training set and using the selected instance as a single-item test set to obtain the sensitivity and the fall-out for different thresholds for the different learning methods (see Tables 2 and 3 for results)

**Table 2.** Machine learning approach used to find GM changes that can eventually discriminate ASD, OCSD and SCHZ. Three different learning methods were applied: the Naive Bayes classifier (Naive), the Support Vector Machines (SVM) and the k-nearest neighbors (KNN).

| GM-KNN | ASD | OCDS | SCHZ |
| --- | --- | --- | --- |
| ASD | / | 23,1% | 25,5% |
| OCSD | 13,3% | / | 31,9% |
| SCHZ | 33,3% | 61.5% | / |

| GM-Naive | ASD | OCDS | SCHZ |
| --- | --- | --- | --- |
| ASD | / | 21.4% | 24.1% |
| OCSD | 30.0% | / | 25.9% |
| SCHZ | 20.0% | 71.4% | / |
| GM-SVM | ASD | OCDS | SCHZ |
| ASD | / | 21.4% | 24.1% |
| OCSD | 30.0% | / | 25.9% |
| SCHZ | 20.0% | 71.4% | / |

**Table 3.**
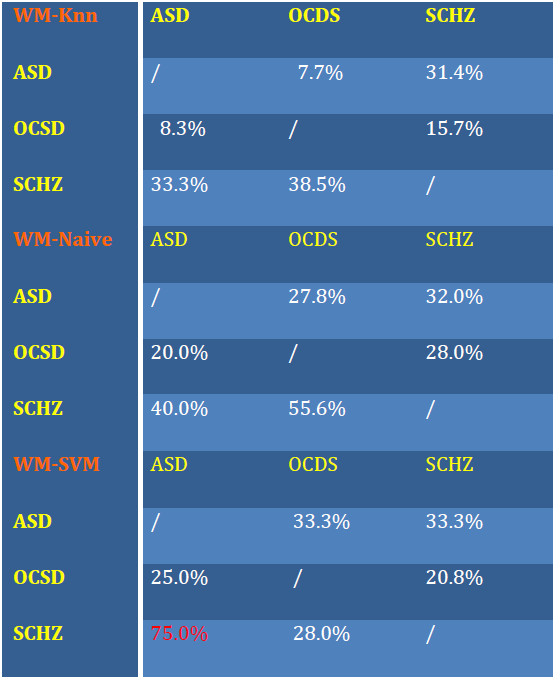
Machine learning approach used to find WM changes that can eventually discriminate ASD, OCSD and SCHZ. Three different learning methods were applied: the Naive Bayes classifier (Naive), the Support Vector Machines (SVM) and the k-nearest neighbors (KNN)

## RESULTS

The GM and WM brain alterations induced by the three spectra considered here can be sub-divided into two clusters as clearly evident in the multidimensional scaling shown in Fig. 1. In both GM and WM we were able to identify a bigger, more homogeneous cluster and a smaller more sparse one. Mapping of the three spectra in the two clusters (see Fig. 2) showed the three spectra to be sparsely distributed in the two clusters. However as shown in Fig. 3 schizophrenia is moderately more present in cluster 1 and ASD spectrum in cluster 2.

**FIG. 1.**
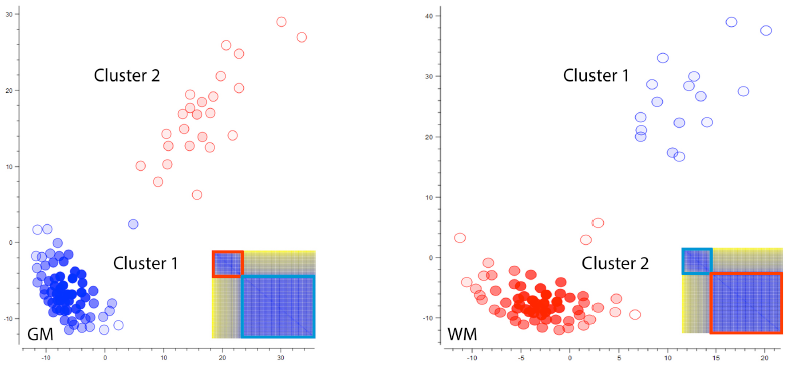
SHOWS THE DISTANCE MATRIX AND THE MULTIDIMENSIONAL SCALING OF THE MA MAPS RELATIVE TO EACH EXPERIMENT HERE INVOLVED. EXPERIMENTS BELONGING TO CLUSTER 1 ARE COLOURED IN BLUE, EXPERIMENTS BELONGING TO CLUSTER 2 ARE COLOURED IN RED.

**FIG. 2.**
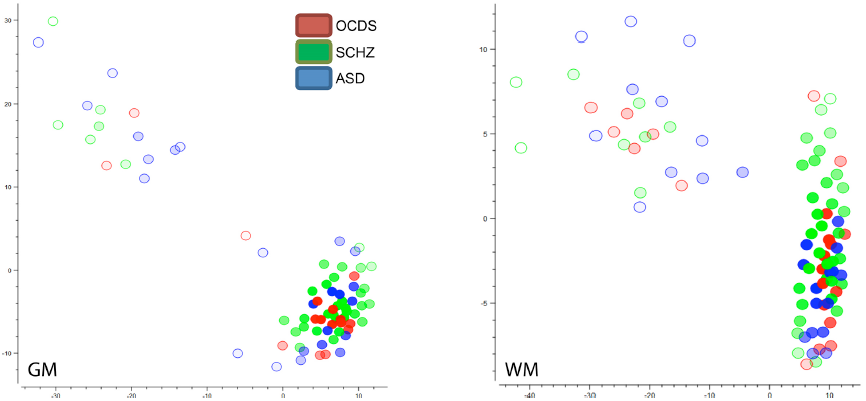
SHOWS THE MULTIDIMENSIONAL SCALING OF THE MA MAPS RELATIVE TO EACH EXPERIMENT HERE INVOLVED. EXPERIMENTS INVOLVING OCDS PATIENTS ARE COLOURED IN RED, EXPERIMENTS INVOLVING SCHZ PATIENTS ARE COLOURED IN GREEN, EXPERIMENTS INVOLVING ASD PATIENTS ARE COLOURED IN BLUE.

**FIG. 3.**
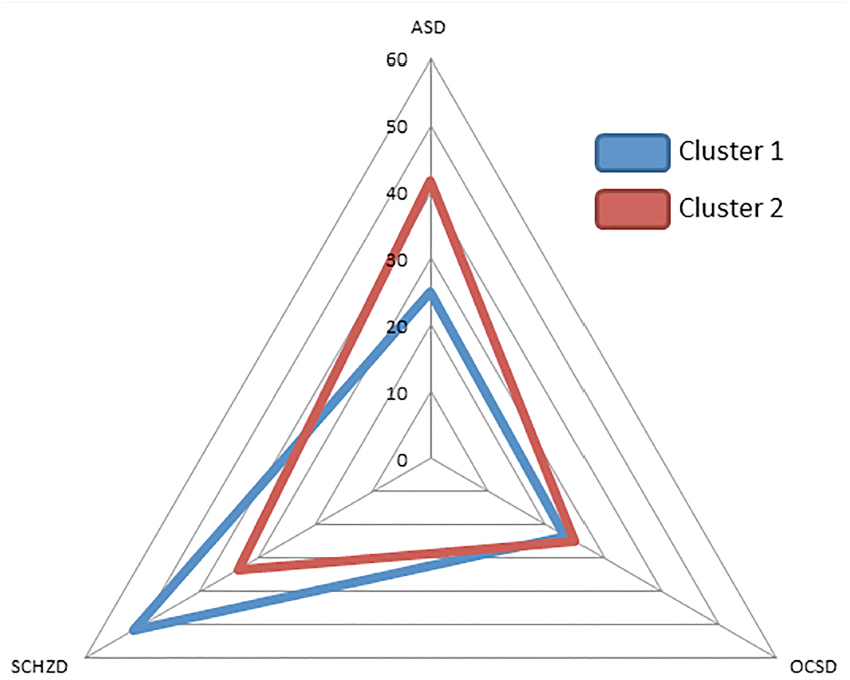
SHOWS THE NUMBER OF EXPERIMENT (IN PERCENT) BELONGING TO EACH OF THE THREE SPECTRUMS THAT ARE CLASSIFIED IN EACH CLUSTER.

The spatial pattern of GM alterations of the two clusters is shown in Fig. 4 and Fig. 5. Cluster 1 (Fig. 4) shows an alteration pattern that closely resembles the “salience detection network” with the anterior insular (AI), anterior cingulate cortex (ACC), dorsolateral prefrontal cortex (DLPFC) and frontopolar brain areas.

**FIG. 4.**
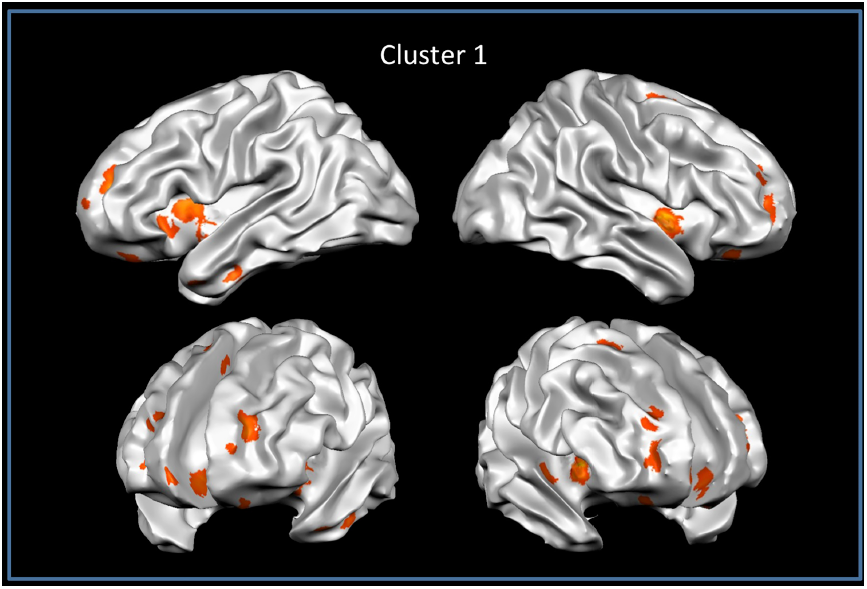
GRAY MATTER ANATOMICAL LIKELIHOOD ESTIMATION (ALE) RESULTS. THIS IMAGE SUMMARIZES THE RESULTS OF ALL THE EXPERIMENTS THAT ARE PARCELLATED TO CLUSTER 1 (ALE MAPS WERE COMPUTED AT A FALSE DISCOVERY RATE CORRECTED THRESHOLD OF P < 0.05, WITH A MINIMUM CLUSTER SIZE OF K > 100 MM^3^ AND VISUALIZED USING BRAINVOYAGER QX).

**FIG. 5.**
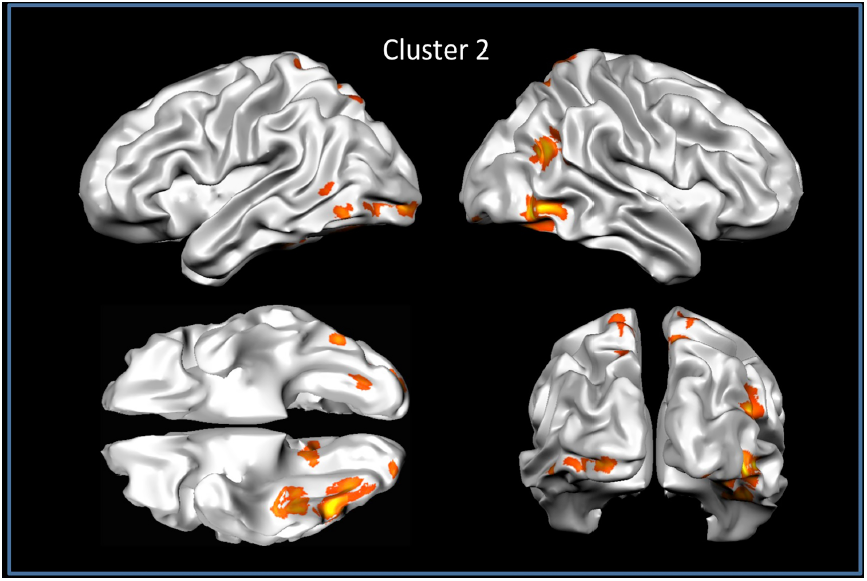
GRAY MATTER ANATOMICAL LIKELIHOOD ESTIMATION (ALE) RESULTS. THIS IMAGE SUMMARIZES THE RESULTS OF ALL THE EXPERIMENTS THAT ARE PARCELLATED TO CLUSTER 2 (ALE MAPS WERE COMPUTED AT A FALSE DISCOVERY RATE CORRECTED THRESHOLD OF P < 0.05, WITH A MINIMUM CLUSTER SIZE OF K > 100 MM^3^ AND VISUALIZED USING BRAINVOYAGER QX).

The second cluster (Fig. 5) shows a more occipital, temporal and parietal spatial pattern with sensorimotor, visual and lingual involvement.

These two patterns partially resemble the “Dual Intertwined Rings Architecture” described by Mesmoudi et al. in a recent paper (Mesmoudi et al., 2013).

### DAMAGE-BASED CLASSIFICATIONS

Data driven classifications of the three spectra on the basis of the pattern of brain alterations reported by each study are shown in Fig. 6 (ROC analysis), and the confusion matrices relative to this classifications in Tables 2 and 3.

**FIG. 6.**
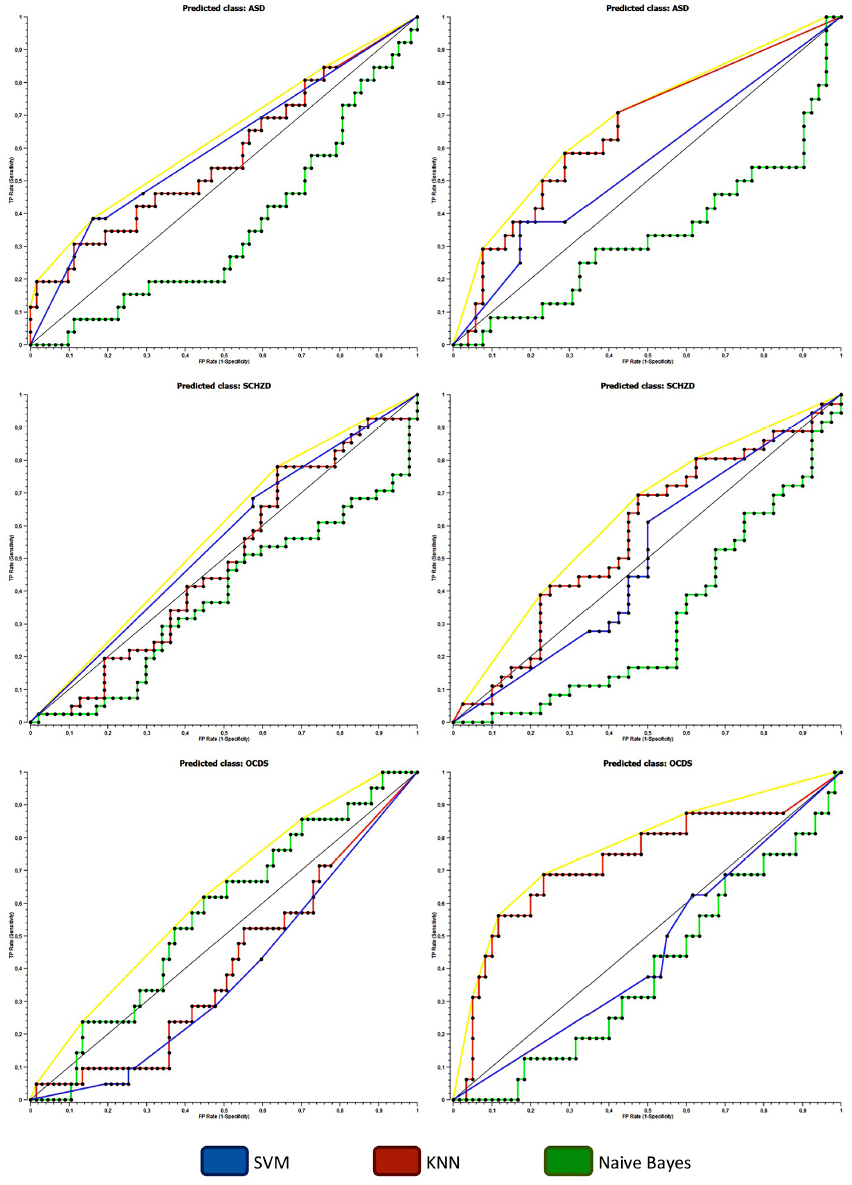
SHOWS THE RECEIVER OPERATING CHARACTERISTIC (ROC) RELATIVE TO EACH CLASSIFICATION APPROACH APPLIED TO GM AND WM DATA.

Using both GM and WM data we were not able to classify the three spectra using a single classifier.

Some classifiers were able to successfully (>chance) classify some pairs of spectra but none were able to correctly classify all of the permutations.

Using GM data all the classifiers correctly classified OCDS vs SCZS. None of the other spectra were correctly classified. Using WM data only one classification was significant: SVM-based Schizophrenia vs ASD.

These results revealed that while OCDS and SCZS showed GM alterations conveying enough differences to be successfully classified, the other pairs of spectra were not differentiable. The relative major distance between OCDS and SCZS was also highlighted by the different presence of the two spectra in clusters 1 and 2 as shown in Fig. 3.

On the whole, the classification approach, together with multidimensional scaling analysis, revealed that the three spectra probably share to a large extent a common set of brain alterations. This fact reduced the classification power of the pattern analysis techniques we employed. The variable part of these alterations was sufficiently informative to classify OCDS vs SCZS only.

### COMMON DAMAGE

Given this result, we then analyzed the commonalities between the GM alterations caused by these three spectra.

An ALE analysis that pooled all the results of the experiments considered in our study (Fig. 7) showed significant GM reductions in the left insular, DLPFC, precentral, inferior frontal, superior temporal and cingulate areas plus the right frontal opercular, postcentral and cingulate areas. GM increases were found in the bilateral AI, inferior temporal and right supramarginal brain areas. A large scale network-based sub-division (see Fig. 8) showed that the networks that are most involved in GM alterations are the salience, premotor, default mode (DMN), basal ganglia-cerebellum and ventral attention network (VAN).

**FIG. 7.**
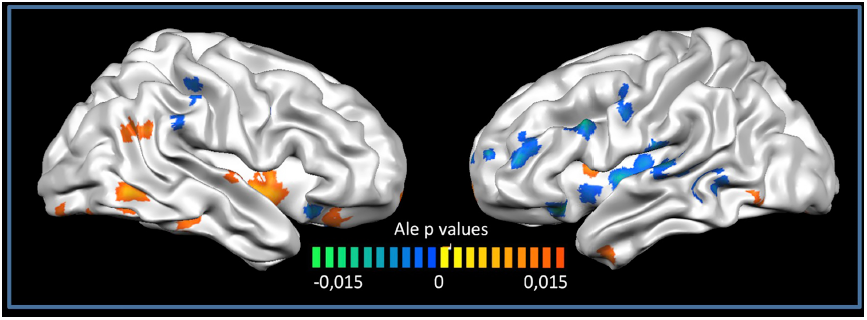
GRAY MATTER ANATOMICAL LIKELIHOOD ESTIMATION (ALE) RESULTS. THIS IMAGE SUMMARIZES THE RESULTS OF ALL THE EXPERIMENTS EMPLOYED IN THIS PAPER. COLORS FROM RED TO YELLOW SHOW GRAY MATTER INCREASES, COLORS FROM BLUE TO GREEN SHOW GRAY MATTER DECREASES (ALE MAPS WERE COMPUTED AT A FALSE DISCOVERY RATE CORRECTED THRESHOLD OF P < 0.05, WITH A MINIMUM CLUSTER SIZE OF K > 100 MM^3^ AND VISUALIZED USING BRAINVOYAGER QX).

**FIG. 8.**
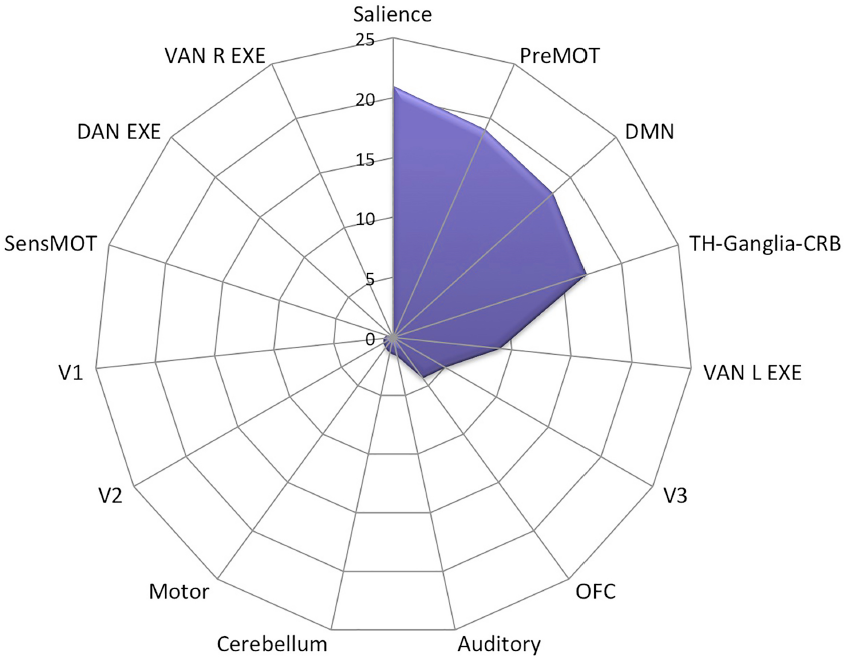
SHOWS THE INVOLVEMENT OF THE LARGE-SCALE BRAIN NETWORKS IN GRAY MATTER ALTERATIONS (EXPRESSED IN PERCENT OF ALTERED VOXELS FOR EACH NETWORKS, ALL EXPERIMENTS TOGETHER).

### DAMAGE NETWORK

The “Damage Network” is shown in Fig. 9. Fig. 10 evidences that the areas with the highest degree are the right supramarginal gyrus (SMG), the thalamus, the amygdalae, the right AI, the putamen, the dorsal ACC (dACC) and the DLPFC. These areas exert the highest number of connections. When damaged, they carry with them a greater number of other damaged-connected brain areas than, for instance, areas with a very low degree (fewer connections) such as the fusiform gyrus. This high degree area thus propagates the damage to a wider set of nodes. These areas are very similar to hub areas in traditional connectivity analysis. In this case however the connections are damage cooccurrences instead of functional, effective or anatomical connections. The interpretation in this case is that the occurrence of damage in a certain area is statistically followed by damage in one or more “damage connected” areas.

**FIG. 9.**
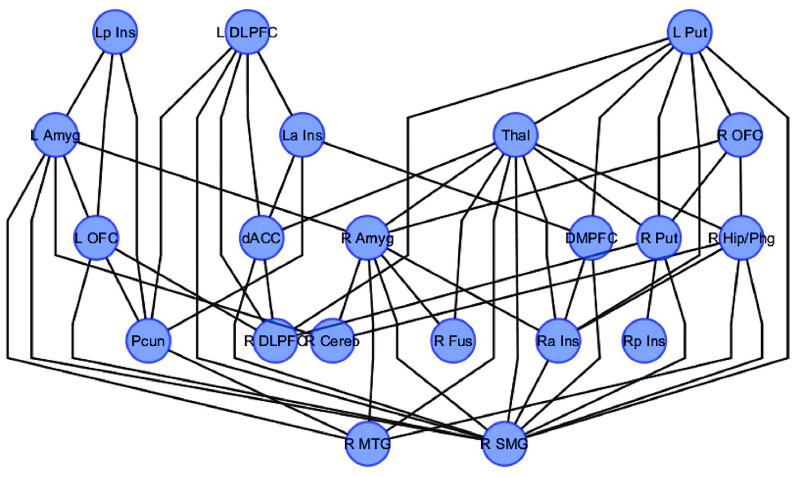
SHOWS A HIERARCHICAL VISUALIZATION OF THE DAMAGE NETWORK. THE NODES OF THE GRAPH ARE PLACED IN HIERARCHICALLY ARRANGED LAYERS SUCH THAT THEEDGES OF THE GRAPH SHOW THE SAME OVERALL ORIENTATION.

**FIG. 10.**
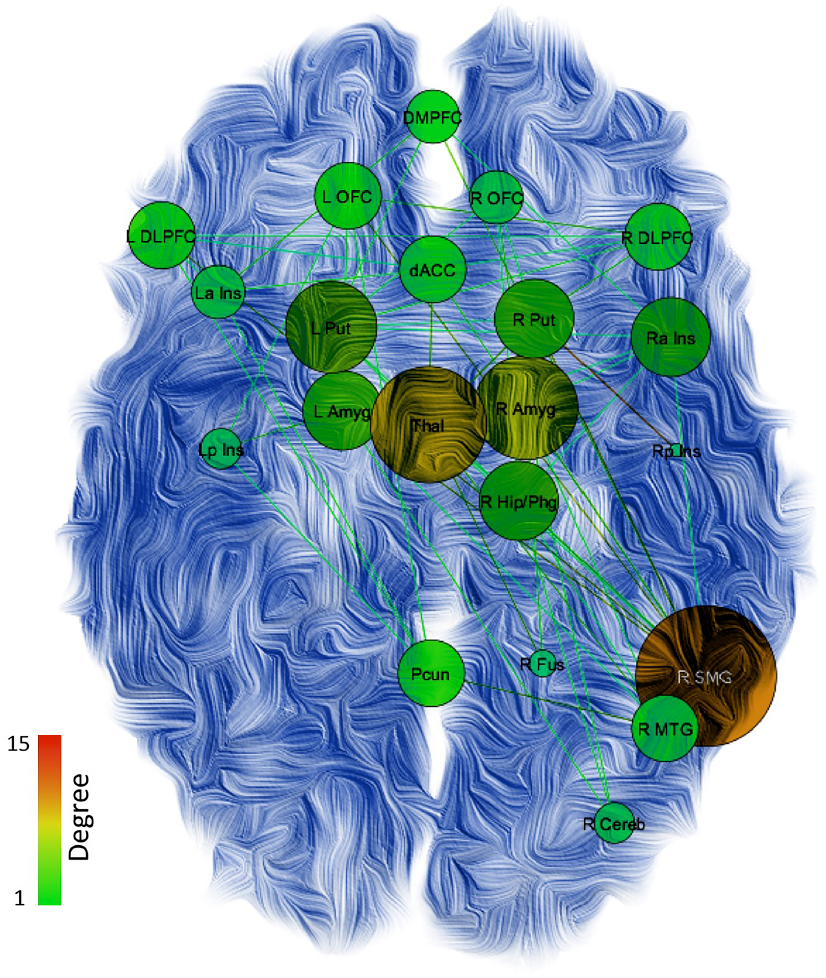
SHOWS THE TOPOLOGICAL ANALYSIS OF THE DAMAGE NETWORK. NODE COLOR AND DIMENSION INDICATES THE DEGREE (SMALLER NODE = LOWER DEGREE; GREEN TO RED = LOWER TO HIGHER VALUES).

Given the relatively large dimension of this network, we investigated the presence of sub-networks in this structure. The k-core decomposition algorithm reported two sub-networks (visualized in Fig. 11); a right network with eight nodes involving the AI, SMG, thalamus, amygdala, hippocampus, putamen and dorsomedial prefrontal cortex (DMPFC); a left cluster with three nodes comprising the DLPFC, AI and dACC.

**FIG. 11.**
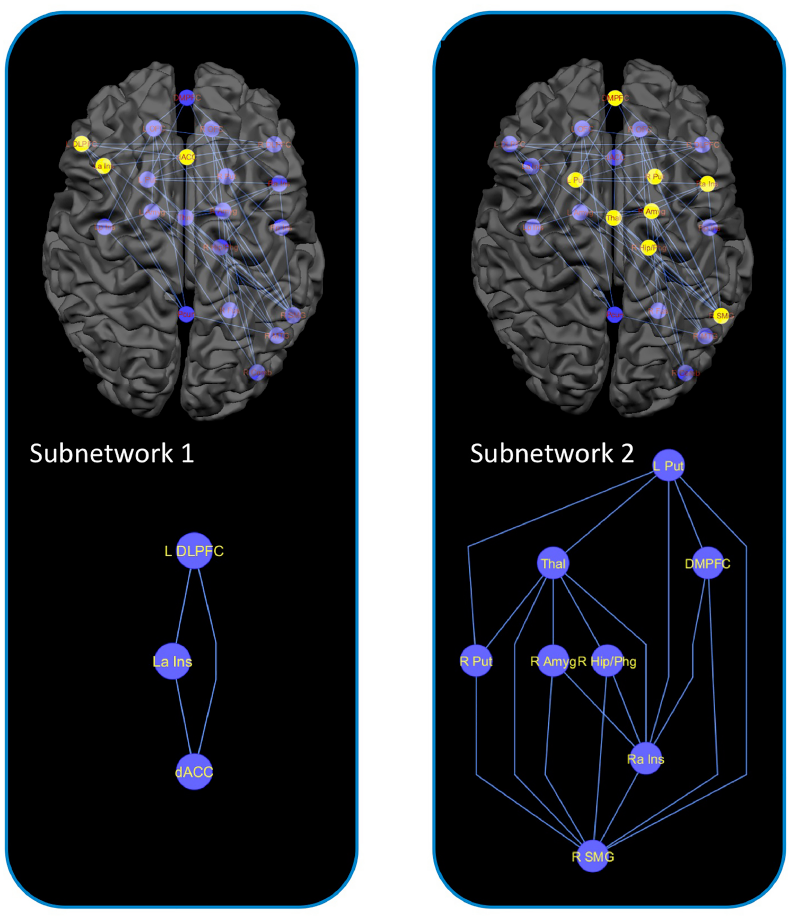
SHOWS THE NETWORK CLUSTERIZATION. THE CLUSTERING ALGORITHM REPORTED TWO CLUSTERS IN OUR DATA: ONE IN THE RIGHT EMISPHERE WITH 8 NODES AND ONE IN THE LEFT HEMISPHERE WITH 3 NODES. LOWER PANEL: HYERARCHICAL REPRESENTATION OF THE LEFT AND RIGHT CLUSTERS. THE NODES OF THE GRAPH ARE PLACED IN HIERARCHICALLY ARRANGED LAYERS SUCH THAT THEEDGES OF THE GRAPH SHOW THE SAME OVERALL ORIENTATION.

## DISCUSSION

In the historical background, linked to Kanner’s (Kanner, 1943) new utilization of Bleuler’s (Bleuler, 1911) autism concept, Ornitz (Ornitz, 1969) observed that the “clinical differences between early infantile autism and its variants and schizophrenia need not be fundamental differences but rather may reflect the influence of maturational level on the overt expression of a single disease process”. Some autistic subjects develop a clinical course that is indistinguishable from schizophrenia, and up to 50% of persons with ASD go on to exhibit psychosis. Furthermore, the gold standard Autism Diagnostic Observation Schedule (ADOS) (Lord et al., 1989) cannot reliably distinguish schizophrenia and child onset schizophrenia from ASD. These clinical observations are supported by several studies demonstrating how the two disorders share some overlaps and similarities in their genetic basis (e.g. GWAS and CNV studies, SHANK 3 variations, DISC 1, dysregulation of CYFIP1, SCN2A, NRXN1 neurexin gene or RELN) (Carroll and Owen, 2009; Guilmatre et al., 2009; Guilmatre et al., 2014; King and Lord, 2011; Pathania et al., 2014; Rapoport et al., 2009), cytoarchitectural organization (e.g. proliferation, migration and lamination defects) (de Lacy and King, 2013), neuropsychological profile (e.g. Theory of mind and mirror neuron function deficits) (Baron-Cohen et al., 1985; Enticott et al., 2008; Lysaker et al., 2011; Rizzolatti and Craighero, 2004; Rizzolatti and Fabbri-Destro, 2010), and neuroimaging patterns (e.g. gray/white matter abnormalities and structural/functional connectivity alterations) (Avino and Hutsler, 2010; Cauda et al., 2014; Cauda et al., 2011; Eastwood and Harrison, 2003; Mueller et al., 2012; Venkataraman et al., 2012; Walterfang et al., 2008).

A comparison of social cognitive functioning in SCZ and high functioning autism shows elevated convergence in particular in SCZ with negative symptoms (Couture et al., 2010; Spek and Wouters, 2010), and neurological soft signs in Asperger syndrome are not different from early-onset psychosis (Mayoral et al., 2010). An overlap of autistic and schizotypal traits in adolescence has been described (Barneveld et al., 2011). Accordingly, a common neurobiological core for SCZS and ASD has been proposed (Bolte et al., 2002; Nylander et al., 2008; Rapoport et al., 2009; Sporn et al., 2004; Starling and Dossetor, 2009; Stone and Iguchi, 2011).

On the other hand, Rutter (Rutter, 1972) drew a clearer separation of autism from schizophrenia in childhood and interestingly in DSM-III (American Psychiatric Association, 1980) autism was incompatible with delusions, hallucinations, loosening of associations, and incoherence (King and Lord, 2011).

OCDS is also clinically related as comorbidity with ASD and SCZS, and a common genetic link among the three spectra has been indicated in proteins such as astrotactins that have a key role in glial-guided neuronal migration during brain development (Lionel et al., 2014). Oxitocin is also higher in OCD SRI-responders and lower in OCD with autistic traits (Humble et al., 2013).

We thus considered addressing the study of ASD, OCDS and SCZS from a neuroimaging perspective (Bastiaansen et al., 2011; Cauda et al., 2011; Jossin and Cooper, 2011; Konstantareas and Hewitt, 2001; Rapoport et al., 2009; Stahlberg et al., 2004; Sverd, 2003; Unenge Hallerback et al., 2012).

Three main findings emerged from our meta-analysis:

1. The three psychiatric spectra we considered do not have their own specific damage. Rather, there is a common injury with some peculiarities typical of each single spectrum. These features are sufficiently informative only for the purpose of correctly classifying the schizophrenic from the autistic spectrum.
2. It is possible to define two different damage clusters. The very reliable first one, constituted by the parietal-temporal-frontal (PTF) ring that includes substantial parts of the salience network, and the second one mainly linked to the auditory-visual, visual-somatic and auditory-somatic (VSA) ring (Mesmoudi et al., 2013). These clusters seem to indicate the presence of damage common to the three pathologies, characterized by cognitive areas (PTF) and by a set of more variable damage that belongs to areas with more functions of input-output (VSA).
3. It is possible, starting from these meta-analytic data, to define a "Damage Network" and to infer a brain substrates hierarchy in the pattern of propagation of the damage.

In our results, as regards the hypothesis of three different spectra, each clearly distinguishable from the others, we expected to find three different clusters of brain alterations. Conversely, starting from data on brain alterations, we found only two clusters: the first (cluster one) more homogeneous and the second (cluster two) more sparse. The three clinical spectra are not specifically related to one of the two clusters, but are distributed in both of the clusters.

In cluster one we found a slight prevalence of SCZS and in cluster two a slight prevalence of ASD. OCDS was more equally distributed between the two clusters.

We found the two clusters to differ in terms of the pattern of brain damage, that in cluster one closely resembles the salience detection network with highly homogeneous findings from a large number of papers, and in cluster two, where the patterns of damage are less homogeneous, relates more to the parietal, inferior temporal and occipital area. Most of the paper belongs to the first cluster. Cluster two shows more posterior brain alterations that are less related to a specific large-scale brain network, but included high order visual, lingual, inferior temporal areas and the precuneus.

These two patterns of damage seem to resemble a recent distinction operated by Mesmoudi et al. (2013) for resting state brain connectivity patterns; in this paper the authors identified a dual intertwined rings architecture of the brain network:

- the parietal-temporal-frontal regions ring (PTF), that relates association cortices specialized in attention, language and working memory to the networks involved in motivation and biological regulation and rhythms and performs “multi-temporal integration that relates past, present and future representations at different temporal scales” (Mesmoudi et al., 2013).
- the visual-somatic-auditory continuous ring (VSA) in which the auditory-visual, visual-somatic and auditory-somatic functions are interspersed between the three monomodal regions, and that performs fast real time multimodal integration of sensorimotor information.

The VSA and PTF rings are intertwined because a subset of associative parietal regions of the PTF ring is at the center of the VSA ring and these regions are connected with other regions of the PTF ring via long-range association tracts passing under the VSA ring.

Intriguingly, our two clusters seem to be part of the two rings, cluster one as part of the PTF and cluster two as part of the VSA.

The PTF ring implements at least four main groups of functions: 1) biological regulation, olfaction, taste and emotion 2) working memory and attention 3) self-referential functions and social cognition 4) language.

Deficit in working memory and attention (lateral-frontal PTF network) has been indicated as a core deficit in schizophrenia strictly related to genetic vulnerability (Bakkour et al., 2014; Tan et al., 2009) and a deficit in executive functions has also been detected in ASD. Impaired social cognition (media-frontoparietal PTF ring) is a link between SCZS and ASD. Language processing by the left temporo-parietal-frontal part of the PTF ring is related to semantic processing, which is mainly altered in ASD but also in schizophrenia. Under-connectivity in frontal-parietal regions has been linked to executive impairments in ASD (Solomon et al., 2009).

In contrast, speech processing that requires real-time processing of visual form perception, motor articulation and auditory perception, is in the VSA ring. This could explain the slight prevalence of ASD in cluster two: in ASD the role of sensoriality (hyper/hyposensoriality as diagnostic criteria) is more relevant than in schizophrenia.

The dual intertwined rings architecture creates three large interfaces between the two rings: the precentral sulcus, the superior temporal sulcus (STS) and the intraparietal sulcus, and places the associative parietal areas BA30 and 40 (which are part of the PTF ring) in the central hole of the VSA ring. The BA39 region is related to the medial part of the PTF ring that is implied in the representations of intentions and social interactions and can thus transform the real-time observation of actions, scenes and events processed in the VSA ring into longer term predictions and interpretations related to intentions and social interactions processed in the PTF ring (Mesmoudi et al., 2013). Gray matter reductions in the superior temporal cortex in schizophrenia have been correlated with the severity of reality distortion symptoms, with delusions and hallucinations (Neckelmann et al., 2006).

The superior temporal cortex has been implicated in face perception and is involved in social cognition, linked to the medial temporal amygdale and hippocampal circuits (Adolphs, 1999).

We tested three different types of classifiers to distinguish the three spectra based on brain damage. Classifiers were not able to find a reliable relationship between brain damage and clinical spectra, albeit with some interesting exceptions. Indeed there is not a single classifier able to classify all three spectra with GM or WM data. The only classification with a clear statistical significance by all three classifiers is the difference between OCD and SCZS. That is to say, with our data it is not possible to reliably distinguish between brain alterations induced by SCZS and ASD and between ASD and OCDS; this may support a link in pathophysiology between these spectra. Thus, even if there is a detectable difference in brain damage between SCZS and OCD, our classification together with the clustering results show a strong convergence between brain alterations in the three spectra. The three spectra show a common “core” of brain damage that is represented in cluster 1 and a set of more variable alterations that are evidenced in cluster 2.

Given these premises, we investigated this common “core” part of brain damage. Fig. 7 shows the “core damage”. GM increases were more frequent in the right hemisphere and reductions in the left hemisphere. The large-scale brain networks damaged most in the three spectra were the salience, premotor, default mode and thalamus-basal ganglia-cerebellum networks. The left ventral attentional, high order visual, auditory and orbitofrontal networks were also involved, but to a lesser extent.

Statistical analysis of the co-activation of damaged areas (that is to say the statistical link between cooccurrence of GM alterations in two or more brain areas) revealed that damage in some GM areas was statistically related to other areas. We defined this concept “Damage Connectivity” and the ensemble of co-damaged areas “Damage Network”. This approach has, for the first time, shed light on the invivo propagation characteristics of brain damage. Our data indicate that not all areas are co-damaged together with other areas and few actually show this type of Damage Connectivity. By plotting the Damage Network we can speculate on the different importance of some areas in the formation of brain damage. Indeed, for example, the right posterior insula is co-damaged (e.g. damage-connected) only with one other area (the right putamen), whereas the right supra-marginal gyrus is co-damaged with eleven other areas. It is thus clear that, if damaged, these two areas may exert a very different role in the Damage Network and may cause a different pattern of propagation of damage in the brain.

Using this innovative method together with network analysis techniques, it is possible to define which areas have the most important role in causing secondary damage (i.e. co-damage in several secondary areas). This data has been defined by calculating the network degree of each damaged area. The areas with the highest network degree are the thalamus, amygdala, putamen and, in the cortical areas, the DLPFC, right AI, right SMG, and the right hippocampus.

Network clustering analysis, performed to determine the presence of any sub-units within the Damage Network, identified two sub-clusters. A left sub-cluster, related to the core areas of the saliency network (AI, DLPFC, dACC) and a more extended right cluster, similarly centered on AI but more widely involving other cortical (right SMG, DMPFC) and sub-cortical areas (amygdala, thalamus, hippocampus, putamen).

At least three neural systems play vital roles in empathy: the mirror neuron system, the affective empathy system located in the AI and midcingulate cortex, and the cognitive empathy system of theory of mind that almost overlaps with the DMN network. The affective empathy system and the cognitive empathy system are linked through the vMPFC (Li et al., 2014).

A link between ASD and SCZS is more closely related to brain areas involved in social cognition and emotion processing, such as the temporal-limbic-frontal pathways related to emotion processing in schizophrenia (Das et al., 2007).

A correlation between the thinning of the fronto-parietal areas and the severity of autistic impairment has been detected and autism and schizophrenia share mirror neuron functional deficits (Enticott et al., 2008; Keller et al., 2011; Rizzolatti and Craighero, 2004; Rizzolatti and Fabbri-Destro, 2010), involved in action understanding (in autism it is possible to understand “what” but not “why”, related to a deficit in chain based mirror mechanism, rather than a hypofunction of the mirror neuron system, in accord with alterations in intra-hemispheric connectivity in ASD) (Boria et al., 2009; Keller et al., 2011).

The AI and the ACC form a salience network, with a key role in ASD and SCZS, that functions to segregate the most relevant among internal and extrapersonal stimuli in order to guide behavior (Menon and Uddin, 2010). The AI is part of a limbic system. It is involved in interoceptive, affective, and empathic processes (Gu et al., 2013) and is part of the salience network integrating external sensory stimuli with internal states and has a relevant role in social processing networks (Anderson et al., 2011; Uddin and Menon, 2009).

The role of the amygdala has been emphasized in ASD and schizophrenia, with under-activation and abnormalities in social processing in both conditions. The left rather than right amygdala has been found to be involved in schizophrenia (Baron-Cohen et al., 2000; Joyal et al., 2003).

The amygdala is linked to the orbitofrontal cortex and superior temporal gyrus in the social brain network and damage to the amygdala is at the root of social impairment. Moreover an early merging neurological insult to the interconnected fronto-amygdala circuit disrupting the ability to flexibly and adaptively orient attention toward self-relevant stimuli might be a primary deficit of ASD. Specifically, the amygdala is responsible, in concert with the vMPFC, for the formation of a priority map of self-relevant events that might be accessible to and modulated by conscious evaluative processing (Zalla and Sperduti, 2013).

The link between ASD and SCZS has a clinical expression in autistic patients with psychotic symptoms that show lower gray matter volume in the right insula than non-psychotic autistic subjects; lower gray matter volume in the right insula is more prominent in schizophrenia than autism (Toal et al., 2009). In adults with ASD, those with a history of psychotic symptoms demonstrated reduced insular volumes, as well as reduced cerebellar volumes (Toal et al., 2009).

Volume alterations in the anterior cingulate cortex, which too is part of the social brain network and is involved in self-perception, social processing, error monitoring and reward-based learning, have also been found in SCZS and ASD (Doyle-Thomas et al., 2013).

The two most prominent brain networks observed during cognitive tasks are the central executive network (CEN), which includes the DLPFC and the posterior parietal cortex, and the default mode network (DMN), which includes the vMPFC and the posterior cingulate cortex: during the performance of cognitively demanding tasks, the CEN typically shows increases in activation, whereas the DMN shows decreased activation.

Lower gray matter volumes in the limbic-striato-thalamic circuitry are common to ASD and SCZS and may partly explain the shared socio-emotional symptoms, while lower gray matter volume in the left putamen (autism) and left fronto-striatal-temporal regions (schizophrenia) appears to distinguish the conditions in terms of gray matter circuitry.

The orbital frontal cortex, in the ventromedial prefrontal region, is thought to play a role in sensory processing, goal directed behavior, adaptive learning and attachment formation and a specific deficit in this area has been correlated with symptom severity in ASD (Ecker et al., 2013).

But the link between the frontal cortex and basal ganglia is the key that joins OCDS, ASD and SCZS. Indeed, the fronto-striatal brain regions are involved in OCD and an orbitofrontal-striatal model has been postulated as an abnormal neural circuit in OCD, although the dorsolateral prefrontal and posterior regions might also be involved in the pathophysiology of OCDS (Cavedini et al., 2002; Nakao et al., 2014). In ASD, caudate volume correlated with repetitive and stereotyped behavior, symptoms related to OCDS (Rojas et al., 2006). Disruption within the basal ganglia loop system is also thought to explain impaired sensorimotor gating, which reflects the ability of an organism to filter out irrelevant stimuli, in both ASD and SCZS. Caudate enlargement, which may be associated with repetitive or self-injurious behavior and a volume loss in the putamen have been reported in ASD (Duerden et al., 2012). The role of the caudate nucleus linked to the orbitofrontal cortex and anterior cingulate cortex has been reported in OCDS (Chamberlain et al., 2005). The cortical-striatal-thalamic loop pathology has been related to a developmental origin (Lipska, 2004), and a developmental syndrome of hippocampal dysfunction has been linked to ASD and SCZS.

The cerebellum has been implicated in SCZS and ASD given its association with learning and cognition (Baribeau and Anagnostou, 2013). Smaller volumes in the cerebellum have been found to be more extensive in autistic subjects with psychosis than in non-psychotic autism (Toal et al., 2009).

## CONCLUSIONS

There is evidence to support the hypothesis that ASD and SCZS are both disorders of cerebral specialization originating in the embryonic period (de Lacy and King, 2013). There is evidence to support the presence of a common neurobiological core, as a shared root of a unique neurodevelopmental tree that could further be categorically divided into different clinical branches. Thus a common ASD-SCZS spectrum could be hypothesized, with a common overlap with OCD spectrum. Further evidence may implicate a common glutamatergic pathomechanism in ASD and SCZS (Gruber et al., 2014) and this could be the link to a common pharmacological role for medication modulating the glutamatergic pathway (Rojas, 2014).

## AKNOWLEDGEMENTS

This study was supported by the Fondazione Carlo Molo, Turin.

## TABLES LEGENDS

Table 1: WM and GM variations with relative numbers of foci for each of the selected psychiatric spectrum.

Table 2: Confusion matrix showing the pattern classification results relative to GM discrimination. Three different learning methods were applied: the Naive Bayes classifier (Naive), the Support Vector Machines (SVM) and the k-nearest neighbors (KNN) to discriminate between ASD, OCSD and SCHZ.

Table 3: Confusion matrix showing the pattern classification results relative to WM discrimination. Three different learning methods were applied: the Naive Bayes classifier (Naive), the Support Vector Machines (SVM) and the k-nearest neighbors (KNN) to discriminate between ASD, OCSD and SCHZ.

